# HIF-1 inactivation empowers HIF-2 to drive hypoxia adaptation in aggressive forms of medulloblastoma

**DOI:** 10.1101/2023.10.17.562750

**Authors:** J. Contenti, Y. Guo, M. Larcher, L. Mirabal-Ortega, M. Rouleau, M. Irondelle, V. Tiroille, A. Mazzu, V. Duranton-Tanneur, F. Pedeutour, I. Ben-Sahra, C. Lago, G. Leva, L. Tiberi, G. Robert, C. Pouponnot, F. Bost, N.M. Mazure

## Abstract

Medulloblastoma (MB) is the most prevalent brain cancer in children. Four subgroups of MB have been identified; of these, Group 3 is the most metastatic. Its genetics and biology remain less clear than the other groups, and it has a poor prognosis and few effective treatments available. Tumor hypoxia and the resulting metabolism are known to be important in the growth and survival of tumors but, to date, have been only minimally explored in MB. Here we show that Group 3 MB tumors do not depend on the canonical transcription factor hypoxia-inducible factor-1α (HIF-1α) to mount an adaptive response to hypoxia. We discovered that HIF-1α is rendered inactive either through post-translational methylation, preventing its nuclear localization specifically in Group 3 MB, or by a low expression that prevents modulation of HIF-target genes. Strikingly, we found that HIF-2 takes over the role of HIF-1 in the nucleus and promotes the activation of hypoxia-dependent anabolic pathways. The exclusion of HIF-1 from the nucleus in Group 3 MB cells enhances the reliance on HIF-2’s transcriptional role, making it a viable target for potential anticancer strategies. By combining pharmacological inhibition of HIF-2α with the use of metformin, a mitochondrial complex I inhibitor to block respiration, we effectively induced Group 3 MB cell death, surpassing the effectiveness observed in Non-Group 3 MB cells. Overall, the unique dependence of MB cells, but not normal cells, on HIF-2-mediated anabolic metabolism presents an appealing therapeutic opportunity for treating Group 3 MB patients with minimal toxicity.

## Group 3 MB are sensitive to oxygen variations and do not express HIF-1 target genes

Medulloblastomas (MBs), like most solid cancers, are subject to continual oxygen variations and have had to develop numerous responses to survive the hypoxic microenvironment^1,2^. The relationship between MB and hypoxia is poorly known. The four different subgroups^3–6^ of medulloblastomas are not similarly located in the cerebellum and access to oxygen is different between the four groups of MB (Fig. 1a). To assess the impact that this oxygen heterogeneity might have on the different MB groups, we analyzed the expression of 19 HIF-1 target genes (*Tf, Ho1*, *Ccl2*, *Oas2*, *Lox*, *Igf2*, *Slc2a1*, *Epo*, *Il2ra*, *Gpnmb*, *bFgf*, *Ak3*, *Hk1*, *Myl3*, *Cxcl16*, *Igfbp2*, *Murc*, *Des* and *Ccl5*) from the Affymetrix Human Gene 1.1 ST array profiling of 763 primary medulloblastoma samples (GSE85217)^7^. Group 4, SHH Group and Wnt Group MBs express these genes most strongly compared to Group 3 MB (Fig.1b). From these data and in order to understand the molecular mechanisms that govern these differences, we focused on the Group 3 MB, the group with a very poor prognosis, using HDMB-03 and D-458 cell lines, and the Non-Group 3 MB, with an intermediate prognosis, using DAOY and ONS-76 cell lines. We cultured these cells in three different oxygen concentrations: 21% O_2_ (normoxia - Nx), in which cells are regularly cultured, 6% O_2_ (physioxia - Phx), the conditions close to physiological conditions of the cerebellum, and 1% O_2_ (hypoxia - Hx), the conditions found in the tumor microenvironment. We observed that Group 3 MB cells were more sensitive to low oxygen conditions, resulting in higher mortality and subsequently lower proliferation than Non-Group 3 MB cells (Fig. 1 c-j). However, all four cell lines expressed HIF-1α and HIF-2α in different profiles, proportions and kinetics (Fig. 1k). Analysis of the different target genes of HIF-1 (*Ca9* and *Mct4/SLC16A3*), HIF-1/HIF-2 (*Ca12*, *Glut1/SLC2A1*, *Ldha*), and HIF-2 (*Oct4/Pou5f1*), and non-target genes of HIFs (*Ldhb* and *MCT1/SLC16A1*) showed that none of the target genes of HIF-1 alone were expressed in Group 3 MB (Suppl. Fig. 1a-h). RNAseq analysis of the four cell lines in Nx *versus* Hx and compared with two Group 3 PDX models, PDX3 and PDX7, confirmed that these models were hierarchically close to the Group 3 cell lines (HDMB-03 and D-458) (Suppl. Fig. 1i). Analysis of genes specific to HIF-1α, HIF-2α and those that could be regulated by either HIF-1α or HIF-2α showed very clearly that Non-Group 3 could induce HIF-1α-specific genes such as *Adm, Angpt1*, *Bnip3l*, *Ca9*, *Lox*, *Slc16a3*, and *Slc2a1* while Group 3 MB as well as the two PDX models representative of Group 3 were unable to (Fig. 1m). In contrast, we observed that HIF-2 specific genes (*MYC*, *Isl2*, *Cdt1*, *PTP4A3*, *MMD*, *Myc*, *Cxcr4*, *GMCL1* and *TRAF4*) were more regulated for in Group 3 MB or model (Fig. 1n). Similar results were found using the R2: Genomics analysis and visualization platform; significantly lower expressions of *Angpt1* and *EPO* were detected in Group 3 MB cells than in Group 4 and/or SHH Group cells (Suppl. Fig. 2). In addition, HIF-2 target genes were shown to have higher expression in Group 3 MB. Similarly, single cell meta data analysis (https://singlecell.broadinstitute.org/) by Manoranjan et al.^8^ using primary patient-derived MB brain tumor-initiating cell (BTIC) lines showed low expression of *Ca9*, *Lox, SLC16A3, ADM* and *Angpt1* in Group 3 MB in a context of high MYC expression compared to the other MB groups whereas *SLC2A1*, *ldha*, *Mpzl1* and *Oct4/Pou5f1*, *Icam4*, the HIF1/2 and HIF-2 target genes, respectively, were expressed in Group 3 MB (Suppl. Fig. 3). These data suggest that Group 3 MB cells have a deficient response to hypoxia and may be due to HIF-1α inability to control its target genes.

**Fig. 1.**
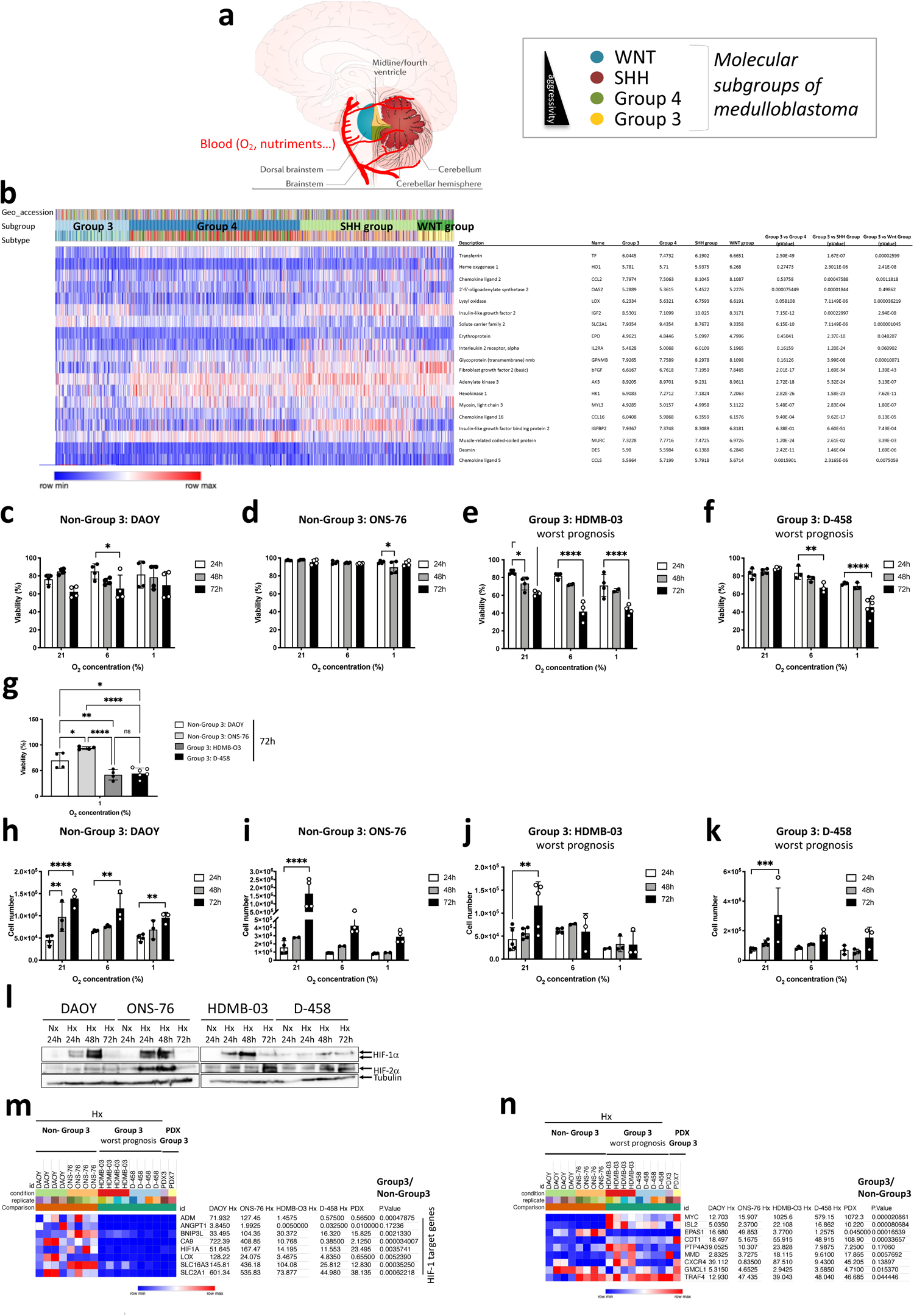
Group 3 MB are sensitive to oxygen variations and do not express HIF-1 target genes. **(a)** Schematic distribution of the different MB subgroups according to the vascular system based on^20^. **(b)** Heatmap of the RNA expression of genes from GSE85217 with 763 primary samples from MB patients from Group 3, Group 4, SHH Group and WNT Group MB^7^. The heatmap focused specifically on target genes of HIF-1. Expression of the genes was compared using Phantasus (v1.19.3). Median expression is represented for each group. p-values were obtained using the Limma differential expression. **c-f:** (**c**) DAOY, (**d**) ONS-76, (**e**) HDMB-03 and (**f**) D-458 cells were incubated in 21%, 6% and 1% O_2_ for 24, 48 and 72 h. Cell viability (%) was measured using an ADAM cell counter. The 2-way ANOVA is representative of at least three independent experiments. * p<0.05, ** p<0.005 and **** p<0.0001. (**g**) Comparison of cell viabilities (%) of DAOY, ONS-76, HDMB-03 and D-458 cells incubated for 72h in 1% O_2_. The 2-way ANOVA is representative of at least three independent experiments. * p<0.05, ** p<0.005 and **** p<0.0001. **h-k:** (**f**) DAOY, (**g**) ONS-76, (**h**) HDMB-03HDMB-03 and (**i**) D-458 cells were seeded at the same density and incubated in 21%, 6% and 1% O_2_ for 24, 48 and 72 h. The 2-way ANOVA is representative of at least three independent experiments. ** p<0.005, *** p<0.0005 and **** p<0.0001. (**l)**, DAOY, ONS-76, HDMB-03, and D-458 cells were incubated in normoxia (Nx, 21% O_2_) for 24h and hypoxia (Hx, 1% O_2_) for 24, 48 and 72 h. Cell lysates were analyzed by immunoblotting for HIF-1α and HIF-2α. Tubulin was used as a loading control. **m-n:** Heatmap of the RNA expression of genes in DAOY, ONS-76, HDMB-03 and D-458 cells exposed to Nx or Hx, compared with PDX3 and PDX7 from Group 3 MB. The heatmap focused specifically on target genes of (**l**) HIF-1 and (**m**) HIF-2. Expression of the genes was compared using Phantasus (v1.19.3). p-values were obtained using the Limma differential expression.

## Post-translational modifications inactivate HIF-1α in Group 3 MB

Immunoblot analysis revealed that HIF-1α from HDMB-03 cells systematically showed a higher molecular weight shift than typical, estimated at about 1 to 2 kDa, while HIF-1α from D-458 cells was mostly absent or otherwise hardly visible (Fig. 2a). As HIF-1α in HDMB-03 was different from classical HIF-1α, we used another anti-HIF-1α antibody to confirm our identification of this transcription factor our observation (Suppl. Fig. 4a). In addition, we verified that the detected bands correspond to HIF-1α using siRNAs (Suppl. Fig. 4b). To better characterize these two HIF-1α from Group 3 MB, we first analyzed their kinetics of stabilization in Nx in the presence of MG132, a proteasome inhibitor (Suppl. Fig. 4c) as well as their stability after reoxygenation (Suppl. Fig. 4e-f). We determined that, overall, MG132 stabilized HIF-1α from Group 3 MB weaklier than Non-Group 3 MB and that HIF-1α was very rapidly degraded in D-458 cells. For HIF-1α in HDMB-03 cells, the band with the highest molecular weight appeared much more stable in Nx even after 10 min of reoxygenation, in contrast to the other cell lines. A 2D gel exploration was possible allowing a finer analysis of HIF-1α in HDMB-03 compared to that in ONS-76. HIF-1α was first stabilized in hypoxia in the presence of Bafilomycin or MG132 or both compounds to block any degradation during sample processing (Suppl. Fig. 4g). Bafilomycin did not stabilize HIF-1α in HDMB-03, suggesting a dysfunction in the mitochondrial uncoupling process^9^, but the use of MG132 in hypoxia increases the stabilization capacity of HIF-1α. Subcellular fractionation was performed on the lysis of both ONS-76 and HDMB-03 cells. Nuclear fractions containing the protein of interest were subjected to 2D electrophoresis. The extracts were either analyzed by immunoblot with an anti-HIF-1α antibody or stained with Coomassie blue and sent to mass spectrophotometry (MS). Under hypoxic conditions, the isoelectric point of the HIF-1α protein was observed at a more basic pH in HDMB-03 compared with ONS-76, strongly supporting a post-translational modification (Fig. 2b). The presence of HIF-1α was confirmed by MS. HDMB-03 cells expressed 6.2 times less HIF-1α protein than ONS-76 (Fig. 2c). The presence of HIF-1β was also characterized. Like HIF-1α, HIF-1β was 15 times less expressed in HDMB-03 cells than in ONS-76 cells, thereby preventing dimerization between the two partners and consequently activation of the target genes. Moreover, we demonstrated that HIF-1α from Group 3 MB cells had little (D-458) or no (HDMB-03) presence in the nucleus (Fig. 2d-e), reinforcing the inability of these cells to activate HIF-1 target genes. HIF-1 cDNA amplification and sequencing revealed no mutation that could explain the different HIF-1α profiles observed in these cells (Suppl. Fig. 5).

**Fig. 2.**
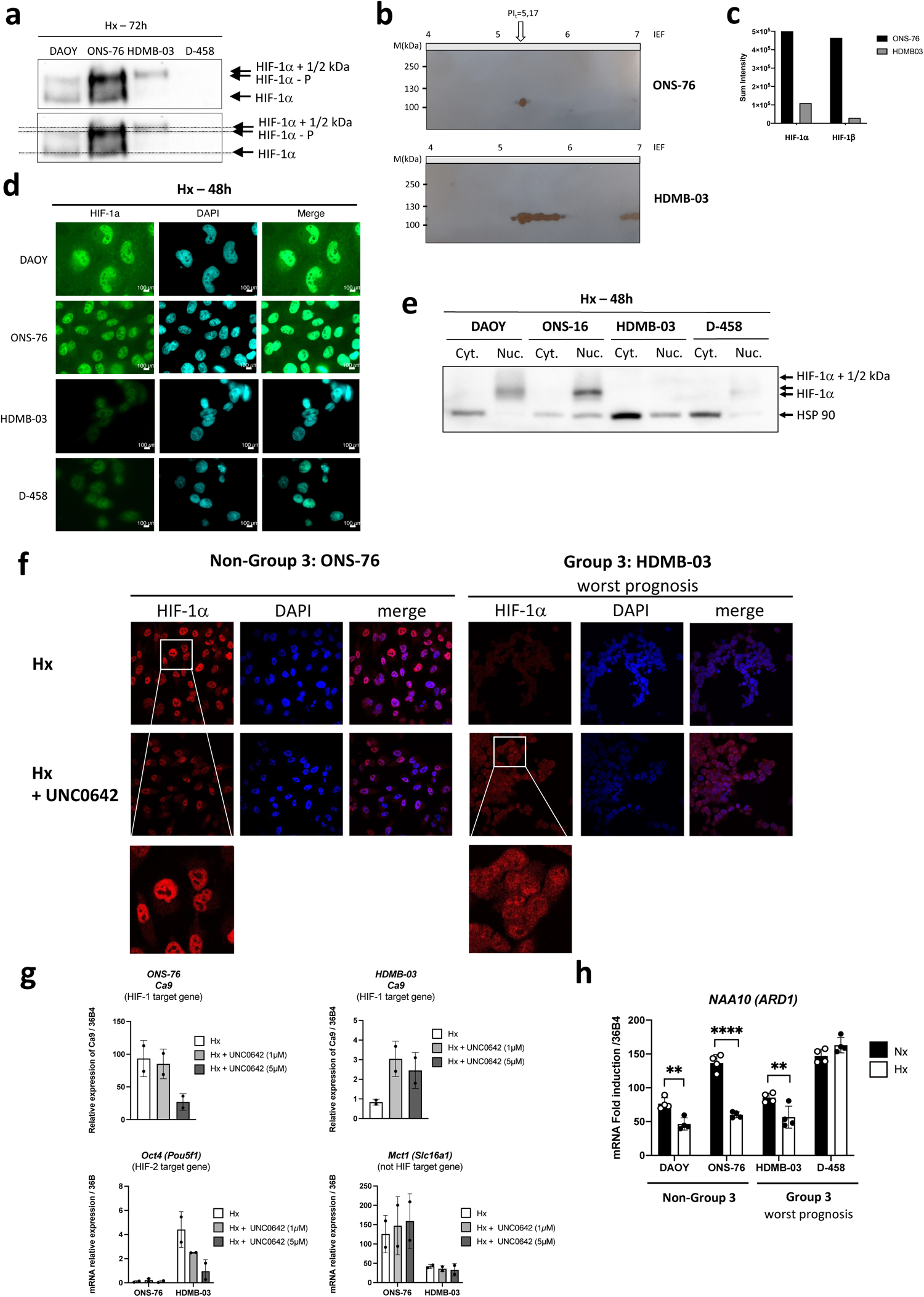
HIF-1α is mutated and transcriptionally inactive in Group 3 MB. **(a)** DAOY, ONS-76, HDMB-03 and D-458 cells were seeded at the same density and incubated in Hx (1% O_2_) for 72h. Cell lysates were analyzed by immunoblotting for HIF-1α. **(b)** HIF-1 protein was induced by hypoxia in ONS-76 and HDMB-03 cell lines. Subcellular fractionation was performed on the lysis of these cells and the nuclear fractions containing the protein of interest were subjected to 2D electrophoresis. The extracts were either analyzed in a Western blot with an anti-HIF-1α antibody or stained with Coomassie blue and sent to mass spectrometry (MS). The horizontal strip at 100 kDa was cut into four equal parts for MS analysis. **(c)** Sum intensity of HIF-1α and HIF-1β in both ONS-76 and HDMB-03 after MS. **(d)** Immunofluorescence labeling and merged images with HIF-1α (in red) and DAPI (in blue) for DAOY, ONS-76, HDMB-03 and D-458 cells incubated for 48h in Hx. **(e)** DAOY, ONS-76, HDMB-03 and D-458 cells were seeded at the same density and incubated in Hx (1% O_2_) for 48h. Subcellular fractionation was used to identify proteins in nuclei and cytoplasm. HSP90 was used as a loading control. **(f)** Immunofluorescence labeling and merged images with HIF-1α (in red) and DAPI (in blue) for ONS-76, and HDMB-03 cells incubated for 48h in Hx in the absence or presence of UNC0642 (5µM). **(g)** Graphic representation of *Ca9*, *Oct4* and *Mct1*mRNA expression in ONS-76 and HDMB-03 cells incubated in hypoxia (Hx - 1% O_2_) for 48h. The 2-way ANOVA is representative of at least three independent experiments. **(h)** Graphic representation of *NAA10* (*Ard1*) mRNA expression in DAOY, ONS-76, HDMB-03 and D-458 cells incubated in normoxia (Nx) for 24h and hypoxia (Hx - 1% O_2_) for 48h. The 2-way ANOVA is representative of four independent experiments. ** p<0.005 and **** p<0.0001.

We thus focused on translational modifications. Phosphorylation of HIF-1α, creating a shift of several kDa, is a central post-translational modification, which regulates its stability under hypoxic conditions (Suppl. Fig. 6a). Overall, *Mekk* (*Map3k1*) and *Erk2* (*Mapk1*) mRNA expression showed no difference in Nx *versus* Hx for either cell line (Suppl. Fig. 6b-d). Only *Erk1* (*Mapk3*) mRNA expression appeared to be induced under hypoxia in ONS-76 (Suppl. 6c). MEKK (MAP3K1) inhibition by U0126 did not block the shift observed on HIF-1α in HDMB-03 in contrast to what we observed in ONS-76, suggesting that this shift is not an additional phosphorylation. Methylation is a post-translational process that results in a 1–2 kDa increase in apparent molecular mass. Methylation can then lead to changes in intracellular distribution, generate protein instability and alter protein-protein interactions ^10^. It has been shown *in vitro* and *in vivo* that HIF-1α can be methylated by G9a/GLP in glioblastoma^11^ (Suppl. Fig. 6f). We therefore hypothesized that HIF-1α expressed in HDMB-03 cells could be methylated. *Ehmt2* (*G9a*) mRNA expression showed no difference between cell lines or between Nx and Hx conditions (Suppl. Figure 6g). On the other hand, while *Ehmt1* (*GPL*) mRNA expression was repressed in Hx in ONS-76, its expression remained unchanged in Hx in HDMB-03, which suggests stronger methylation by *Ehmt1* (*GPL*) (Suppl. Fig. 6h). Moreover, *Ruvbl2* (*Reptin*) mRNA expression was also less repressed in Hx in HDMB-03 cells compared with ONS-76 cells (Suppl. Fig. 6i). Using UNC0642, a small molecule inhibitor specific for EHMT2/1, we showed that demethylated HIF-1α regained its ability to be stabilized (Suppl. Fig. 6j), to accumulate in the nucleus (Fig. 2f) and to activate its target gene (*Ca9*) (Fig. 2g), strongly supporting that HIF-1α from the HDMB-03 is methylated in Hx. Oct4 expression was decreased, suggesting that HIF-2α needs to be methylated to be active while MCT1 expression was not affected by the presence of UNC0642. Although HIF-1α was better stabilized in the presence of UNC0642, we did not observe a decrease in the shift. K32 and/or K391 methylation of HIF-1α seemed not to be involved as PFI-2, a potent inhibitor of SETD7, did not improve HIF-1α stabilization (Suppl. Figure 6k-n). To reinforce these findings, we compared the expression of *Ehmt1*, *Ehmt2* and *Ruvbl2* in the Non-Group 3 and Group 3 MBs using the GSE85217 microarray of 763 MB patient tissues. Only *Ruvbl2* was significantly more expressed in Group 3 MB. As Group 3 had 3 subgroups (alpha, beta, gamma), we looked at the expression of these 3 genes according to the subgroups. It was in Group 3 gamma, the most aggressive group, that *Ehmt2* and *Ruvbl2* genes were significantly more expressed. With the hypothesis that overexpression of EHMT2 and RUVBL2 would lead to methylation and/or transcriptional inactivity of HIF-1, we compared the expression of HIF-1 target genes in the 3 subtypes. We found a lower expression of 14 HIF-1 target genes (*Cacna1s*, *Ccgng2*, *Ccng2*, *Cited2*, *Flt1p1*, *Hk2*, *Igfbp2*, *Lox*, *Pkm1/2*, *Slc2a3*, *Sl2a1*, *Tgfb3*, *Tpi1* and *Ttn*) that was more pronounced in Group 3 gamma, reinforcing the potential involvement of the EHMT1/2/RUVBL2 pathway in the control of HIF-1 (Suppl. Figure 7). The genes thus impacted are particularly involved in cellular metabolism (Suppl. Figure 7c).

We then investigated the reason of the absence of HIF-1α in D-458 by studying the full process of stabilization of the protein under hypoxic conditions. pVHL binds and degrades HIF-1α only when HIF-1α is hydroxylated by the prolyl hydroxylases (PHDs) in the presence of oxygen (Suppl. Fig. 8a). In the absence of oxygen, the PHDs are inhibited, allowing stabilization of HIF-1α. However, a feedback loop is established and *phd*s will certainly be induced to ensure rapid degradation under reoxygenation. An increase in the quantity of PHDs could also counteract their low activity. Auto-regulation in hypoxia occurs through increased expression of the genes *phd2* and *phd3*, but not *phd1* since these genes are themselves HIF-targets. Non-Group 3 cells showed consistent regulation of expression of all three *phds*, in contrast to Group 3 cells (Suppl. Fig. 8b-d). *phd1* expression, already high in Nx, was not modulated in hypoxia in D-458 cells. *Phd2* was not modulated in Hx in D-458 cells and *phd3* was non-existent in both Group 3 MB cell lines. We also checked if *Vhl* had mutations in the different lineages, but we did not find any mutation (data not shown) and the expression of *Vhl* mRNA did not show any discordance between Groups (Suppl. Fig. 8e). However, while DAOY, ONS-76 and HDMB-03 cells showed statistically lower proteosomal activity in Hx, D-458 cells showed a higher trend under these same conditions (Suppl. Fig. 8f-g). Finally, since acetylation of K532 by the acetyltransferase ARD1 promotes interaction with pVHL and thus degradation of HIF-1α in the proteasome, we checked *Ard1* (*Naa10*) mRNA expression. While *Ard1* expression was repressed in Hx in DAOY, ONS-76 and HDMB-03 cells, again, we found that ARD1 expression, already very high in Nx, was not altered in Hx in D-458 cells (Fig. 2h). However, invalidation of *Ard1* by siRNA did not stabilize HIF-1α (data not shown). Interestingly, significantly higher expressions of *Egln1, Egln3* and *Naa10* were detected in Group 3 MB than in the Non-Group 3 using the R2: Genomics analysis and visualization platform (Suppl. Figure 8h-i). Egln3 and Egln1 mRNA expression were also found to be significantly upregulated using the GSE85217 microarray (Suppl. Figure 8k).

Taken together, these results strongly suggest that Group 3 MB cells have nonfunctional HIF-1 resulting from post-translational modifications of HIF-1α, methylation in the HDMB-03 cells and high destabilization in D-458 cells, and thus that the response of Group 3 MB cells to the hypoxic microenvironment relies essentially on HIF-2.

## PT2385 sensitizes Group 3 MB to cell death *in vitro* and decreases glycolysis

We then considered that the use of an HIF-2α inhibitor could specifically target the metabolism of Group 3 MB. We chose PT2385, one of the first specific HIF-2α inhibitors identified, currently used in clinical trials for kidney cancer and glioblastoma. The lowest concentration of PT2385 (1µM) decreased viability specifically in Group 3 MB (Fig. 3a). As expected, in Hx (Fig. 3b-c) and Phx (data not shown), PT2385 reduced glycolytic capacity by 35.3% and 38.2% in HDMB-03 and D-458 cells, respectively, in somewhat similar proportions to those observed with 2-DG used as a control (Suppl. Fig. 9a). The repressive effect of PT2385 was not observed in Nx, in which HDMB-03 showed even better glycolytic and respiratory capacities with PT2385 (Suppl. Fig. 10). No deleterious effect was observed in Non-Group 3 MB (Fig. 3d-e). Finally, PT2385 had no effect on respiration in Hx in either Group 3 MB (Fig. 3f-g) or Non-Group 3 MB (Fig. 3 h-i). It may, however, have increased breathing capacity in Nx, as observed in HDMB-03 (Suppl. 10). To completely collapse the energy intake of the Group 3 MB, we explored blocking the respiration with Metformin (Metf), a well-known inhibitor of the complex 1 of the respiratory chain and oxygen consumption. Metf clearly decreased the proliferation of all cells in Hx while specifically targeting only Group 3 MB viability both in Phx and Hx (Fig. 3j-k). The addition of Metf did block respiration in Group 3 MB cells but only partially decreased respiration in Non-Group 3 MB cells (Suppl. Fig. 9b). A mirror effect on glycolysis was observed, as Metformin increased glycolysis overall in Group 3 MB to compensate for the lack of energy due to the blockage of respiration but this compensatory effect was barely observed in the Non-Group 3 MB (Suppl. Fig. 9).

**Fig. 3.**
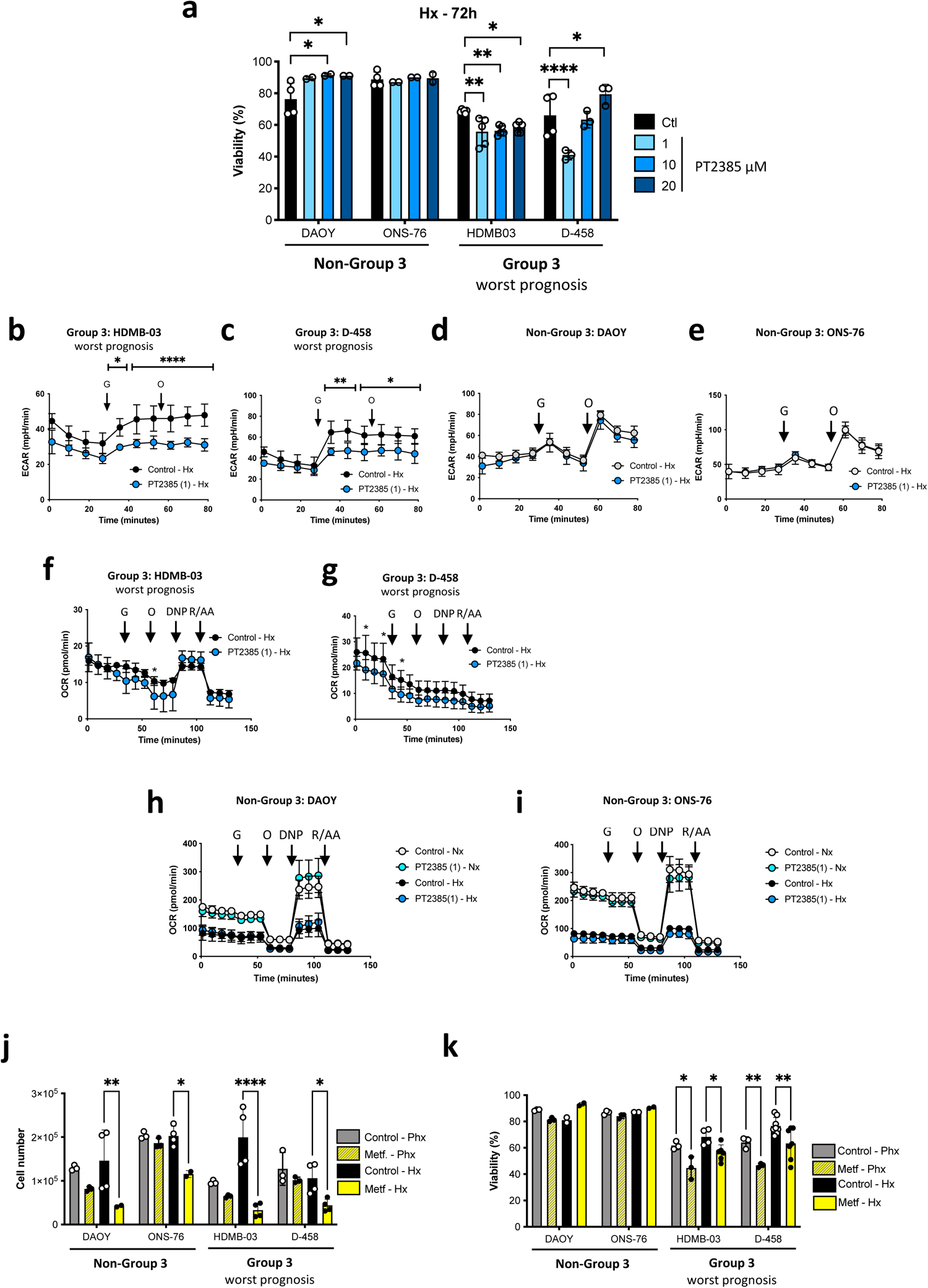
PT2385 sensitizes Group 3MB viability i*n vitro* and affects glycolysis. **(a)** DAOY, ONS-76, HDMB-03, and D-458 cells were incubated in Hx for 72h in the absence (Control) or presence of PT2385 at 1-, 10-, and 20-µM. Cell viability (%) was measured using an ADAM cell counter. The graphs are representative of at least three independent experiments carried out in octuplicate. * p<0.05, ** p<0.005 and **** p<0.0001. **b-e:** Glycolysis in the absence (Control) or presence of PT2385 of **(b)** HDMB-O3, **(c)** D-458, **(d)** DAOY and **(e)** ONS-76 cells in Hx (1% O_2_) for 24h. ECAR was evaluated with the XF24 analyzer. Cells were deprived of glucose for 1h, then glucose (G) and oligomycin (O) were injected at the indicated times. The graphs are representative of at least three independent experiments carried out in octuplicate. * p<0.05, ** p<0.005 and **** p<0.0001. **f-i:** Respiratory control of **(f)** HDMB-O3, **(g)** D-458, **(h)** DAOY and **(i)** ONS-76 cells, cultured for 24h in Hx (1% O_2_) in the absence (Control) or presence of PT2385. OCR was measured in real time with the XF24 analyzer. Cells were deprived of glucose for 1h, then glucose (G), oligomycin (O), FCCP (F), and Rotenone + Antimycin A (R/A) were injected at the indicated times. The graphs are representative of at least three independent experiments carried out in octuplicate. * p<0.05. **(j)** DAOY, ONS-76, HDMB-03 and D-458 cells were incubated in Phx (6% O_2_) or Hx (1% O_2_) for 72h in the absence (Control) or presence of Metformin (Metf). Cell number was measured using an ADAM cell counter. The graphs are representative of at least three independent experiments carried out in octuplicate. * p<0.05, ** p<0.005 and **** p<0.0001. **(k)** DAOY, ONS-76, HDMB-03 and D-458 cells were incubated in Phx (6% O_2_) or Hx (1% O_2_) for 72h in the absence (Control) or presence of Metformin (Metf). Cell viability (%) was measured using an ADAM cell counter. The graphs are representative of at least three independent experiments carried out in octuplicate. * p<0.05 and ** p<0.005.

It appears that the significant biological effects of PT2385 and Metf together is limited to Group 3 MB suggesting that their combination could be of therapeutical interest.

## PT2385/Metf combo decreases Group 3 MB cell viability and induces major metabolic modifications

We thus looked at the impact of these compounds, alone or in combo, on the stabilization of HIF-1/2α. PT2385 alone slightly decreased the stabilization of HIF-2α only in D-458 cells without affecting HIF-1α stabilization (Suppl. Fig. 11a). However, the combination of the two compounds strongly affected HIF-1α and HIF-2α stability (Suppl. Fig. 11b). This resulted in decreased expression of the genes *Ca9*, *Ca12*, *Glut1*, and *Ldha* (Suppl. Fig. 11c-g) but not of *Ldhb* (Suppl. Figure 11h). Subsequently, dual PT2385 and Metformin treatment clearly decreased proliferation in all cell lines (Suppl. Fig. 12a-b) and decreased viability in particular in Group 3 MB in Phx and Hx (Fig. 4a-b). SiRNAs directed against HIF-1α in the presence of PT2385 had no effect on cell viability in the Group 3 MB, whereas nearly 40% of the cells in the Non-Group 3 MB died, unable to depend on any HIF-α sub-unit (Fig. 4c). In addition, siRNA directed against HIF-2α showed no effect on cell viability in the Non-Group 3 MB, whereas it slightly reinforced the action of PT2385/Metf in the Group 3 MB. Glycolysis as well as respiration were entirely collapsed in Group 3 MB but only partially in Non-Group 3 MB (Suppl. Fig. 12c-j).

**Fig. 4.**
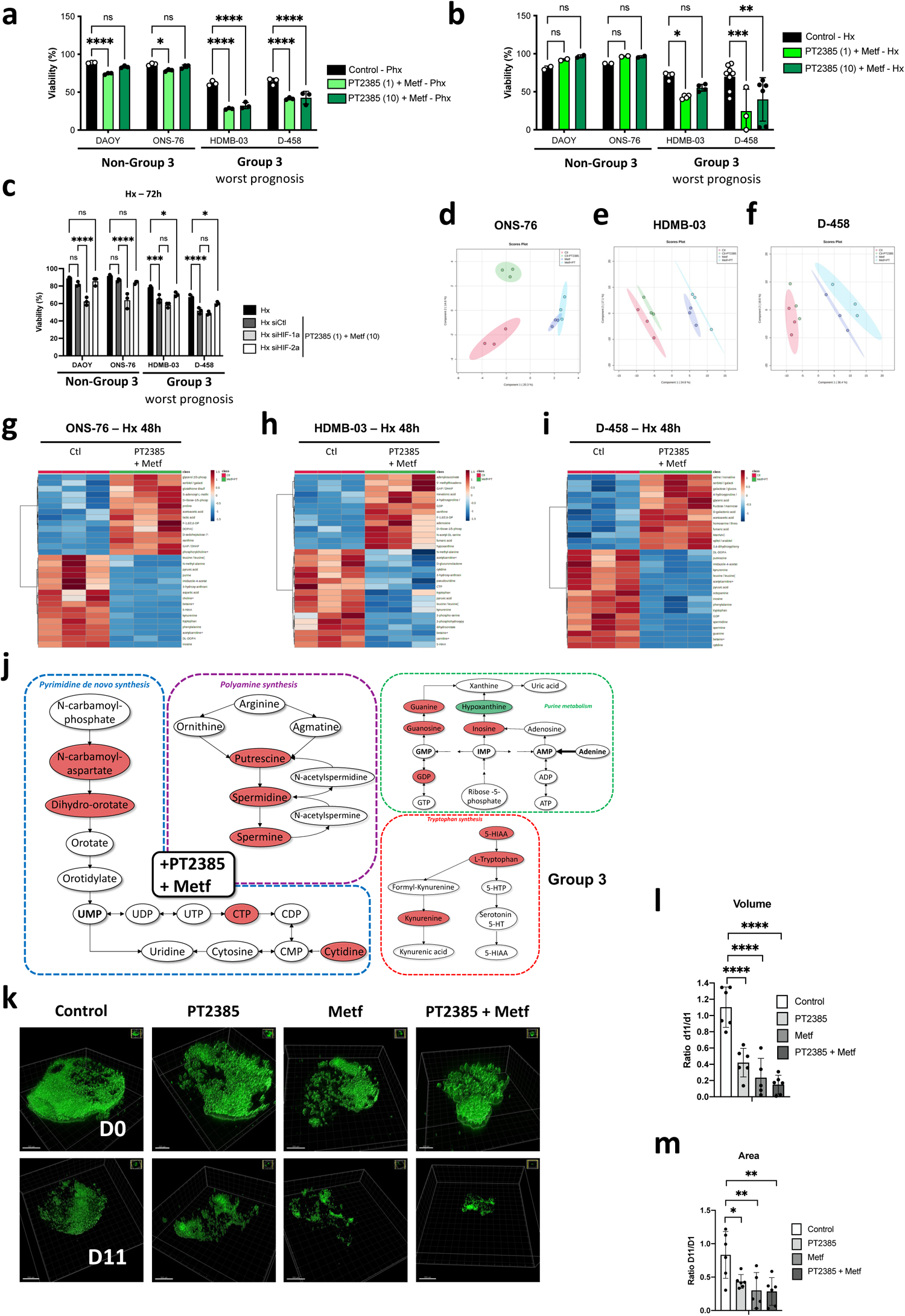
PT2385/Metf combo is effective *in vitro* and induces Group 3 MB cell death. **a-b:** DAOY, ONS-76, HDMB-03 and D-458 cells were incubated in (**a**) Phx or (**b**) Hx for 72h in the absence (Control) or presence of PT2385 (1- or 10-µM) and Metformin (Metf, 10mM). Cell viability (%) was measured using an ADAM cell counter. The 2-way ANOVA is representative of at least three independent experiments. * p<0.05, ** p<0.005, *** p<0.0005 and **** p<0.0001. **(c)** DAOY, ONS-76, HDMB-03 and D-458 cells were transfected with a control siRNA (siCtl) or an siRNA specific to HIF-1α (siHIF-1α) for 24h. Cells were then incubated in hypoxia (Hx) for 72h. Cell viability (%) was measured using an ADAM cell counter. The 2-way ANOVA is representative of three independent experiments. * p<0.05, *** p<0.0005 and **** p<0.0001. **d-f:** Principal component analysis (PCA) scores plot in positive mode based on samples of (**d**) ONS-76, (**e**) HDMB-03 and (**f**) D-458 cells, subjected to 24h of Hx (1% O_2_), comparing cells in the absence (Ctl) and presence of PT2385 (1µM), Metf (10mM) or both. **g-i:** Steady-state metabolite profile of (**g**) ONS-76, (**h**) HDMB-03 and (**i**) D-458 cells subjected to hypoxia (Hx) for 48h in the absence (Ctl), or presence of PT2385+Metf. Intracellular metabolites from three independent samples per condition were profiled by LC/MS-MS, and those significantly altered in treated cells, relative to control cells, are shown as row-normalized heatmaps ranked according to fold-change log_2_ (treated/untreated). **(j)** Schematic representation of the different major metabolic pathways impacted by PT2385 + Metf treatment in Group 3 MB versus ONS-76 (Non-Group 3 MB). **(k)** Three-dimensional structures at day 0 (D0) and day 11 (D11) obtained from confocal image series using IMARIS software; scale bars = 500 µm. **l-m:** Quantification of (**l**) cell volume, and (**m**) cell area at day 1 and day 11. A ratio was calculated from day 11 to day 1. The 2-way ANOVA is representative of at least five different organoids. * p<0.05, ** p<0.005 and **** p<0.0001.

Multivariate analysis of metabolomic profiling of ONS-76, HDMB-03 and D-458 untreated or treated with PT2385, Metf or both in Hx, using principal component analysis (Fig. 4d-e) and hierarchical clustering (Fig. 4g-i and Suppl. Fig. 13a-f) revealed common or different profiles depending on the treatments used. We highlighted clear changes in metabolites related to polyamine pathways and purine and pyrimidine synthesis in the presence of PT2385 alone in Group 3MB compared to ONS-76 (Suppl. Fig. 13g). Similar changes were observed with Metf but with a stronger impact than PT2385 suggesting an interesting yet unexploited role for Metf in MB (Suppl. Fig. 13h). In addition, the tryptophan synthesis pathway was decreased (Fig. 4 j). Interestingly, the use of the combo had a real additive effect blocking more strongly the four metabolic pathways previously described (Fig. 4j and Suppl. Fig. 13g-h).

Together, these results strongly suggest that the combination of PT2385 and Metf would (i) simultaneously block glycolysis and respiration in the Group 3 MB and (ii) act directly on nucleotide synthesis. These different actions represent a promising therapeutic approach.

## Blocking HIF-2 pathway in combination with Metf induces apoptosis of Group 3 tumor organoids

Finally, we sought to validate the drug responses identified in 2D on an organoid-derived model of human Group 3 MB in which cancer cells are GFP/Venus positive ^12^. Human Group 3 MB organoids were generated with MYC and Otx2 overexpression, to recapitulate the gene alteration/expression of Group 3 MB patients. We examined the effects of the two drugs, PT2385 and Metf, alone and in combination. The organoids were maintained for 11 days and treated every 4 days with the different compounds. We observed significant antitumor activity of Metf and PT2385+Metf at day 4 (Suppl. Fig. 14a and b). Venus fluorescence disappeared almost completely by day 11, showing a nearly total disappearance of the tumor cells with the combo. However, the effect of PT2385 seemed weaker, its anti-tumoral action appearing later (D8). Confocal image stacks of the tumoral Venus cells were reconstructed to 3D isosurfaces (Imaris) (Fig. 4k). The volume (Fig. 4l), and area (Fig. 4m) showed significant decreases compared to the control. Finally, control cells showed less cleaved caspase 3 staining than cells in the presence of the different treatments (Suppl. Fig. 14c). Super imposition of images double stained with green (Venus) and red (cleaved caspase 3) fluophores showed that tumor cells expressed both proteins, strongly suggesting that cells died of apoptosis in the presence of the combo but also in the presence of the compounds alone.

These findings suggest that the drug responses observed *in vitro* can be recapitulated when assayed in an *in vivo* context with the human Group 3 MB tumor organoids.

## Discussion

Here we show for the first time that cell lines belonging to Group 3 MB, the most aggressive and metastatic group, exhibit post-translational modifications of HIF-1α that lead to functional inactivation of HIF-1. The HIF-2 isoform is then the only one left to respond to different oxygen variations. This makes it interesting to compare Group 3 MB with ccRCCs expressing only HIF-2, a particular type of aggressive kidney cancer. The activity of HIF-1α, rather antitumorigenic in this specific cancer, is strongly decreased due to chromosomal deletions^13^ or HAF activity^14^ leaving HIF-2 to fully exert its pro-tumor activity. The dichotomy between HIF-1 and HIF-2 observed in ccRCCs was until now unique in the tumor field. What about medulloblastomas and especially Group 3 MB? In the Non-Group 3, the presence of a functional HIF-1 coupled to HIF-2 seems to protect the tumor cells from cell death and none of the tested treatments seemed to affect them. However, what our results suggest is more that HIF-1 introduces a survival capacity in a hostile environment like hypoxia. Therefore, the absence of HIF-1α would not appear to benefit the tumor cell. However, HIF-1 acting alone does in fact give the cell an advantage. Group 3 MB has been characterized by high expression of the MYC oncogene that enables rapid and aggressive tumor development^15^. HIF-2α is a formidable promoter of MYC activity while HIF-1α tends to inhibit it. Moreover, MYC has been shown to regulate the HIF-2α expression^16^. Pei et al. showed that MYC overexpression in stem cells had strong morphological homologies and expression profiles comparable to those of Group 3 cells^15^. Their studies suggest that MYC overexpression is a major oncogenic event for Group 3 MB: it induces tumorigenesis and then allows its maintenance and progression. However, HIF-2 is a formidable player in maintaining stem cells in an undifferentiated state *via* genes like *Oct4*, *Sox* and *Nanog* ^17,18^. In physioxia or hypoxia where Oct4 would be induced, the cells would then have a strong stem cell potential even though these cells would be “adult”. Since hypoxia occurs as early as the embryonic state^19^, we can propose that HIF-2 may be the primary origin of oncogenic evolution of Group 3 MB. If the uniqueness of HIF-2 is an extraordinary advantage for the Group 3 MB tumor cell, then it becomes a perfect target to block MB tumor growth. We used PT2385, which is in phase II for ccRCC (https://clinicaltrials.gov/ct2/show/NCT03108066) but also for glioblastoma (https://clinicaltrials.gov/ct2/show/NCT03216499). Very promising results have been obtained, suggesting similar results would be possible for Group 3 MB patients. Other more efficient inhibitors identified recently, such as PT2399 or PT2977, could certainly provide stronger therapeutic responses. HIF-2 inhibition could then not only partially block metabolism but also affect particular stem cells. However, unlike ccRCC, it was the combination of PT2385 with Metf that had a very strong impact on cell growth and death. Here we have proven the concept that the best targeted therapy comes from a better molecular knowledge. It seems clear that this “hypoxic” approach will reveal other potential therapeutic targets and allow new drug combinations to be found, to provide the most specific and personalized therapeutic response possible.

## Acknowledgments

This work was supported by a grant from La Fondation Flavien, the Fondation ARC pour la recherche sur le Cancer and La Ligue contre le Cancer. Y. Guo is supported by the Chinese Ministry of Research. I. Ben-Sahra is supported by the NIH R00-CA194192 and LAM Foundation grants. F. Bost and N.M. Mazure are CNRS investigators. Metabolomics services were performed by the Metabolomics Core Facility at Robert H. Lurie Comprehensive Cancer Center of Northwestern University and the Beth Israel Deaconess Medical Center Mass Spectrometry Facility of Harvard Medical School. Armenise, AIRC, CARITRO and EMBO to L.T.

We sincerely thank the GIS-IBISA multi-sites platform Microscopy Imagery Côte d’Azur (MICA), and particularly the imaging site of C3M (INSERM U1065) supported by INSERM, Cancéropôle PACA, Conseil Régional, Conseil Départemental, and IBISA.

## Conflict of interest statement

The authors declare no potential conflicts of interest.

---

**Suppl. Figure 1. a-h:** Graphic representation of (**a**) *Ca9*, (**b**) *Mct4*, (**c**) *Ca12*, (**d**) *Glut1*, (**e**) *Ldha*, (**f**) *Oct4*, (**g**) *Mct1* and (**h**) *Ldhb* mRNA expression in DAOY, ONS-76, HDMB-03, and D-458 cells incubated in Nx for 24h and Hx for 24, 48 and 72 h.

The 2-way ANOVA is representative of at least three independent experiments. * p<0.05, ** p<0.005, *** p<0.0005 and **** p<0.0001. **(i)** Heatmap of the RNA expression of genes in DAOY, ONS-76, HDMB-03 and D-458 cells exposed to Nx or Hx, compared with PDX3 and PDX7 from Group 3 MB.

**Suppl. Figure 2. a-h**, Boxplot distributions of gene expressions of (**a**) *HIF-1α*, (**b**) *Angpt1*, (**c**) *Epo*, (**d**) *HIF-2α/EPAS*, (**e**) *Isl2*, (**f**) *Myc*, (**g**) *Vegfa* and (**h**) *Ldha* in Group 3, Group 4, SHH Group and WNT Group MB cells using the R2: Genomics analysis and visualization platform.

**Suppl. Figure 3.** (**a)** tSNE of 3725 cells colored by GSVA enrichment score for the Wnt subgroup gene signature: Group 3 (red), Group 4 (blue) and Wnt Group (green)^8^.

**b**-**j:** Cells colored by classification for enrichment of Wnt subgroup signature. Dark red cells surpassed the 5% cutoff and are considered significantly enriched for the gene signature. Gray cells did not pass the threshold. (**b**) *Ca9*, (**c**) *Lox*, (**d**) *Glut1* (*Slc2a1*), (**e**) *Ldha*, (**f**) *Mpzl1*, (**g**) *Oct4* (*Pou5f1*), (**h**) *Icam4*, (**i**) *Mct1* (*Slc16a1*), (**j**) *ldhb* and (**k**) *Myc*.

**(i)** HIF-1, HIF-1/2 and HIF-2 percentage of expression in the different subgroups (WNT, GR4 and GR3).

**Suppl. Figure 4. (a)** ONS-76 and HDMB-03 cell lysates were analyzed by immunoblotting for HIF-1α from Invitrogen (PA5-85494) and Tubulin was used as a loading control. (**b)** DAOY, ONS-76, HDMB-03 and D-458 cells were transfected with control siRNA (-) and a pool of siHIF-1α ((+) siHIF1pool). Cell lysates were analyzed by immunoblotting for HIF-1α, HIF-2α, and Tubulin was used as a loading control. **(c)** DAOY, ONS-76, HDMB-03 and D-458 cells were seeded at the same density and incubated in Nx (21% O_2_) for 18h. MG132 (10µM) was added for 1, 2, 4 or 6 h. Cell lysates were analyzed by immunoblotting for HIF-1α. Tubulin was used as a loading control. Bottom panel, Quantification of HIF-1α protein levels. The 2-way ANOVA is representative of at least three independent experiments. * p<0.05, *** p<0.0005 and **** p<0.0001. (**d)** DAOY, ONS-76, HDMB-03, and D-458 cells were incubated in hypoxia (Hx) for 48h and reoxygenated for 1, 2, 4, 6, and 10 min (’). Cell lysates were analyzed by immunoblotting for HIF-1α. Tubulin was used as a loading control. (**e)** Quantitative analysis of the total HIF-1α bands compared to tubulin. (**f)** Quantitative analysis of the upper and lower HIF-1α bands compared to tubulin. (**g)** HDM-03 and ONS-76 cells were seeded at the same density and incubated in Hx (1% O_2_) for 48h. Bafilomycin (Baf) or MG132 or both (+MG132+Baf) were added 6h prior to lysis. Subcellular fractionation was used to identify proteins in nuclei and cytoplasm. Tubulin was used as a loading control.

**Suppl. Figure 5. (a)** Summary of primers used for RT-PCR. (**b)** Representative experiments of RT-PCR of HIF-1α cDNA. (**c)** Alignment of the cDNA of HIF-1α (NCBI Reference Sequence: NC_000014.9) with the cDNA of HIF-1α from DAOY, ONS-76, HDMAB03 and D-458 cells. (**d)** Alignment of the HIF-1α protein sequences from HIF-1α control (sp|Q16665|HIF1A_HUMAN Hypoxia-inducible factor 1-alpha), DAOY, ONS-76, HDMB-03 and D-458.

**Suppl. Figure 6. (a)** Molecular signal affecting the phosphorylation of HIF-1α.

**b-d:** Graphic representation of (**b**) *Mekk (MAP3K1)*, (**c**) *Erk1 (Mapk3)*, and (**d**) *Erk2 (Mapk1)* mRNA expression in ONS-76 and HDMB-03 cells incubated in normoxia (Nx) or hypoxia (Hx - 1% O_2_) for 24h. The 2-way ANOVA is representative of at least three independent experiments. * p<0.05. **e**, ONS-76 and HDMB-03 cells were seeded at the same density and incubated in Hx (1% O_2_) for 48h in the absence (-) or presence of (1 and 10 µM) U0126. ONS-76 and HDMB-03 cell lysates were analyzed by immunoblotting for HIF-1α and Tubulin was used as a loading control. **(f)** Molecular signal affecting the methylation of HIF-1α through EHMT1/2. **g-i:** Graphic representation of (**g**) *Ehmt2 (G9a)*, (**h**) *Ehmt1 (Gpl)*, and (**i**) *Ruvbl2 (Reptin)* mRNA expression in ONS-76 and HDMB-03 cells incubated in Nx or Hx for 24h. The 2-way ANOVA is representative of at least three independent experiments. * p<0.05 and ** p<0.005. (**j)** ONS-76 and HDMB-03 cells were seeded at the same density and incubated in Hx for 48h in the absence (-) or presence (+) of UNC0642 (5µM). ONS-76 and HDMB-03 cell lysates were analyzed by immunoblotting for HIF-1α and Tubulin was used as a loading control. **(k)** Molecular signal affecting the phosphorylation of HIF-1α through SETD7. **l-m:** Graphic representation of (**l**) *Setd7 (Set7/9)* and (**m**) *Kdm1a (Lsd1)* mRNA expression in ONS-76 and HDMB-03 cells incubated in Nx or Hx for 24h. The 2-way ANOVA is representative of at least three independent experiments. *** p<0.0005. (**n)** ONS-76 and HDMB-03 cells were seeded at the same density and incubated in Hx (1% O_2_) for 48h in the absence (-) or presence (+) of PFI-2 (2 nM (+) and 2µM (++)). ONS-76 and HDMB-03 cell lysates were analyzed by immunoblotting for HIF-1α and Tubulin was used as a loading control.

**Suppl. Figure 7. a-b:** Violin distributions of gene expressions of *Ehmt1, Ehmt2* and *Ruvbl2* in (**a**) Non-Group 3 *versus* Group 3 MB and (**b**) subgroups (alpha, beta and gamma) of Group 3 MB. (**c**) Top - Violin distributions of HIF-1 target gene expressions in subgroups (alpha, beta and gamma) of Group 3. Bottom - KEGG Fold enrichment from ShinyGo 0.77. **(a-c)** The 2-way ANOVA is representative of at least three independent experiments. * p<0.05, ** p<0.005, *** p<0.0005 and **** p<0.0001. (**d**) Network representation of KEGG from ShinyGo 0.77.

**Suppl. Figure 8. (a)** Molecular signal affecting the hydroxylation and acetylation of HIF-1α. **b-e**, Graphic representation of (**b**) *Phd1 (Egln2)*, (**c**) *Phd2 (Egln1)*, (**d**) *Phd3 (Egln3)* and (**e**) *Vhl* mRNA expression in ONS-76 and HDMB-03 cells incubated in normoxia (Nx) for 24h or hypoxia (Hx - 1% O_2_) for 48h. The 2-way ANOVA is representative of at least three independent experiments. * p<0.05, *** p<0.0005 and **** p<0.0001. **f-g:** DAOY, ONS-76, HDMB-03 and D-458 cells were seeded at the same density and incubated in Nx or Hx for 48h. (**f**) Proteasome activity in the cells, measured in pMol/well. The 2-way ANOVA is representative of two independent experiments. * p<0.05 and ** p<0.005. (**g**) Graph showing the ratio of the proteasome activity obtained in Nx *versus* Hx for each of the cells. **h-j:** Boxplot distributions of gene expressions of (**h**) *egln1* (*phd2*), (**i**) *egln3* (*phd3*) and (**j**) *naa10* (*ard1*). (**k**) Violin distributions of gene expressions of *Egln3 (Phd3), Egln1 (Phd2)* and *vhl* in Non-Group 3 *versus* Group 3 MB.

**Suppl. Figure 9. (a)** ECAR of DAOY (first line), ONS-76 (second line), HDMB-03 (third line) and D-458 (fourth line) cells cultured in Nx (21% O_2_ – first column), Phx (6% O_2_ – second column) and Hx (1% O_2_ – third column) in the presence of Metformin (Metf) or 2-DG for 24 h was evaluated with the XF24 analyzer. Cells were deprived of glucose for 1 h, then glucose (G) and oligomycin (O) were injected. The graphs are representative of at least three independent experiments carried out in octuplicate. Yellow star (*) represents the statistical differences between Metf and control and red (*) star between 2-DG and control. The 2-way ANOVA is representative of at least three independent experiments. * p<0.05, ** p<0.005, *** p< 0.001 and ****p<0.0001.

**(b)** OCR of DAOY (first line), ONS-76 (second line), HDMB-03 (third line) and D-458 (fourth line) cells cultured in Nx (21% O_2_ – first column), Phx (6% O_2_ – second column) and Hx (1% O_2_ – third column) in the presence of Metformin (Metf) or 2-DG for 24 h was evaluated with the XF24 analyzer. Cells were deprived of glucose for 1h, then glucose (G), oligomycin (O), DNP, and Rotenone + Antimycin A (R/A) were injected at the indicated times. The graphs are representative of at least three independent experiments carried out in octuplicate. Yellow star (*) represents the statistical differences between Metf and control. The 2-way ANOVA is representative of at least three independent experiments. * p<0.05, ** p<0.005, *** p< 0.001 and ****p<0.0001.

**Suppl. Figure 10. a-b,** ECAR of (**a**) HDMB-03 and (**b**) D-458 cells in Nx in the absence (Control) or presence of PT2385 (1µM) for 24h was evaluated with the XF24 analyzer. Cells were deprived of glucose for 1h, then glucose (G) and oligomycin (O) were injected at the indicated times. The graphs are representative of at least three independent experiments carried out in octuplicate. **c-d,** Respiratory control of **(c)** HDMB-03 and **(d)** D-458 cells. OCR was measured in real time with the XF24 analyzer. Cells were cultured for 24h in Nx in the absence (Control) or presence of PT2385 (1µM). Cells were deprived of glucose for 1h, then glucose (G), oligomycin (O), DNP, and Rotenone + Antimycin A (R/A) were injected at the indicated times. The graphs are representative of at least three independent experiments carried out in octuplicate. * p<0.05, ** p<0.005 and ****p<0.0001.

**Suppl. Figure 11. (a)** DAOY, ONS-76, HDMB-03 and D-458 cells were incubated in hypoxia (Hx - 1% O_2_) for 72 h in the absence (-) or presence (+) of PT2385 (1µM). Cell lysates were analyzed by immunoblotting for HIF-1α and HIF-2α. Tubulin was used as a loading control. (**b)** DAOY, ONS-76, HDMB-03 and D-458 cells were incubated in Hx for 72 h in the absence (-) or presence (+) of PT2385 (1µM) and Metformin (Metf – 10mM). Cell lysates were analyzed by immunoblotting for HIF-1α and HIF-2α. Tubulin was used as a loading control. **c-h:** Graphic representation of (**c**) *Ca9*, (**d**) *Ca12*, (**e**) *Glut1 (Slc2A1)*, (**f**) *Oct4 (Pou5f1)*, (**g**) *Ldha* and (**h**) *Ldhb* mRNA expression in DAOY, ONS-76, HDMB-03 and D-458 cells incubated in Hx for 48h in the absence (Control) or presence of PT2385 (1µM) or the presence of both PT2385 (1µM) and Metformin (Metf – 10mM). The 2-way ANOVA is representative of at least three independent experiments. Non-significant (ns), * p<0.05, ** p<0.005, *** p< 0.001 and ****p<0.0001.

**Suppl. Figure 12. a-b**, DAOY, ONS-76, HDMB-03 and D-458 cells were incubated in **(a)** physioxia (Phx) and **(b)** hypoxia (Hx – **b**) for 72h in the absence (Control) or presence of PT2385 (1µM or 10µM) and Metformin (Metf – 10mM). Cell number was measured using an ADAM cell counter. The 2-way ANOVA is representative of at least three independent experiments. ** p<0.005, *** p< 0.001 and ****p<0.0001. **c-f:** ECAR of (**c**) DAOY, (**d**) ONS-76, (**e**) HDMB-03 and (**f**) D-458 cells cultured in Hx in the presence of Metformin (Metf – 10mM), PT2385 (1µM) or both for 24h was evaluated with the XF24 analyzer. Cells were deprived of glucose for 1h, then glucose (G) and oligomycin (O) were injected. Yellow star (*) represents the statistical differences between Metf and control and green (*) star between PT2385+Metf and control. The 2-way ANOVA is representative of at least three independent experiments. * p<0.05, *** p< 0.001 and ****p<0.0001. **g-j:** OCR of (**g**) DAOY, (**h**) ONS-76, (**i**) HDMB-03 and (**j**) D-458 cells cultured in Hx in the presence of Metformin (Metf – 10 mM), PT2385 (1µM) or both for 24 h was evaluated with the XF24 analyzer. Cells were deprived of glucose for 1h, then glucose (G), oligomycin (O), DNP, and Rotenone + Antimycin A (R/A) were injected at the indicated times. Yellow star (*) represents the statistical differences between Metf and control and green (*) star between PT2385+Metf and control. The 2-way ANOVA is representative of at least three independent experiments. * p<0.05, ** p<0.005, *** p< 0.001 and ****p<0.0001.

**Suppl. Figure 13. a-f,** Steady-state metabolite profile of (**a-b**) ONS-76, (**c-d**) HDMB-03 and (**e-f**) D-458 cells submitted to hypoxia (Hx) for 48h in the absence (Ctl), or presence of PT2385 or Metf. Intracellular metabolites from three independent samples per condition were profiled by LC/MS-MS, and those significantly altered in treated cells, relative to control cells, are shown as row-normalized heatmaps ranked according to fold-change log_2_ (treated/untreated). **(g)** Schematic representation of the different major metabolic pathways impacted by PT2385 treatment in Group 3 MB *versus* ONS-76. **h,** Schematic representation of the different major metabolic pathways impacted by Metf treatment in Group 3 MB *versus* ONS-76.

**Suppl. Figure 14. (a)**Validation of drug response in human Group 3 MB tumor organoids. Brightfield and fluorescence images of cerebellar organoids at day 0-, day 4-, day 6-, day 8-, day 11- and day 14-of treatment, electroporated with pBase + pPBMYC + pPBOtx2 + pPBVenus. **(b)** Quantification of the mean (raw integrated density/area using Fiji) for each condition from day 4 (D4) to day 14 (D14). The ordinary two-way ANOVA is representative of at least three independent organoids. * p<0.05, ** p<0.005 and *** p=0.0006. **(c)** Superimposition images where the presence of both fluors, Venus (tumor cells) and red (cleaved-caspase 3) is shown as a third color (yellow - merge). Human Group 3 MB tumor organoids have been treated in the absence (Ctl) or presence of PT2385, Metf or the combo. The orthogonal view is used to show virtual cross sections – one plotted along the x-axis and the other plotted along the y-axis; scale bars = 100 µm.

## STAR Methods

### Cell Culture

DAOY (from ATCC - HTB-186^TM^) and ONS-76 (from Dr. F. Di Cunto (University of Torino - Italy)) cells were grown in Dulbecco’s Modified Eagle’s Medium (DMEM) (Gibco-BRL, Courtaboeuf, France) supplemented with 10% fetal bovine serum with penicillin G (50 U/mL) and streptomycin sulfate (50g/mL) whereas HDMB-03 (from DSMZ - ACC740) and D-458 cells, provided by Dr. C. Pouponnot (Institut Curie - France), were grown with the same medium supplemented with 20% fetal bovine serum. A BugBok workstation (Ruskinn Technology Biotrace International Plc, The Science Park Bridgend, UK) set at 6% oxygen, 94% nitrogen and 5% carbon dioxide was used for physioxic conditions. A Whitley H35 hypoxystation anaerobic workstation (Don Whitley Scientific, West Yorkshire, UK) set at 1% oxygen, 94% nitrogen and 5% carbon dioxide were used for hypoxic conditions.

### Patient-derived xenografts

MB patient-derived xenograft (PDX) models were developed from primary tumor samples of previously untreated patients, implanted into the neck fat pad of Nude mice^21^. In all cases, primary human brain tumor specimens were obtained under written informed consent approved by the Internal Review Board of the Necker Sick Children’s Hospital, Paris, France. The protocol also complied with internationally established 3R principles, in accordance with the UKCCCR guidelines. Once established, PDX models were maintained by serial passages in Nude mice. PDX3 and PDX7 correspond to IC-MB-PDX-1 and ICN-MB-PDX-7 respectively.

### Pharmacological Inhibitors and Chemicals

Cells were incubated with 10 mM Metformin (Metf) to block mitochondrial Complex I, 2-DG (10mM) to block glycolysis and PT2385 (1- and 10-mM) to inhibit HIF-2α activity. Rotenone, antimycin A, oligomycin, and 2,4-Dinitrophenol (DNP) were from Sigma, (St. Louis, MI, USA).

### RNA interference

The 21-nucleotide RNAs were chemically synthesized (Eurogentec, Seraing, Belgium) and previously described (PMID: **16585195**). The siRNA sequences, all validated, were as follows: siCtl (forward) 5’-CCU-ACA-UCC-CGA-UCG-AUG-AUG-TT-3’, siHIF-1α (forward) 5’-CUG-AUG-ACC-AGC-AAC-UUGATT-3’, siHIF-2α (forward) 5’-CAG-CAU-CUU-UGA-UAG-CAG-UTT-3’.

### PCR analysis

Total RNA was extracted with the RNeasy Mini Kit (QIAGEN, Hilden, Germany). The amount of RNA was evaluated with a NanoDrop™ spectrophotometer (ThermoFisher Scientific, Waltham, MA USA). One μg of total RNA was used for reverse transcription, using the QuantiTect Reverse Transcription kit (QIAGEN, Hilden, Germany), with oligo (dT)_15_ to prime first-strand synthesis. Full-length HIF-1α cDNAs of approximately 3.5 kb were amplified by RT-PCR different primers shown in Suppl. Figure 4a. The HIF-1α cDNAs were amplified and sequenced.

### Quantitative real-time PCR analysis

Total RNA was extracted with the RNeasy Mini Kit (QIAGEN, Hilden, Germany). The amount of RNA was evaluated with a NanoDrop™ spectrophotometer (ThermoFisher Scientific, Waltham, MA USA). One μg of total RNA was used for reverse transcription, using the QuantiTect Reverse Transcription kit (QIAGEN, Hilden, Germany), with oligo (dT)_15_ to prime first-strand synthesis. SYBR master mix plus (Eurogentec, Liege, Belgium) and specific oligonucleotides (Sigma Aldrich) were used for qPCR. Primer sequences used were: *Ca9* (forward: 5’-CCGAGCGACGCAGCCTTTGA −3’; reverse: 5’-GGCTCCAGTCTCGGCTACCT-3’), Ca12 (forward: 5’-CTGCCAGCAACAAGTCAG-3’; reverse: 5’-ATATTCAGCGGTCCTCTC-3’), *Glut1* (forward: 5’-CTTCACTGTCGTGTCGCTGT −3’; reverse: 5’-TGAAGAGTTCAGCCACGATG-3’), *Oct4* (forward: 5’-TGGAGTTTGTGCCAGGGTTT-3’; reverse: 5’-CTGTGTCCCAGGCTTCTTT-3’), *Ldha* (forward: 5’-AGCCCGATTCCGTTACCT-3’; reverse: 5’-CACCAGCAACATTCATTCCA-3’), *Ldhb* (forward: 5’-GATGGATTTTGGGGGAACAT-3’; reverse: 5’-AACACCTGCCACATTCACAC-3’), *Mct4* (forward: 5’-ATTGGCCTGGTGCTGCTGATG-3’; reverse: 5’-CGAGTCTGCAGGAGGCTTGTG-3’); *Mct1* (forward: 5’-CACCGTACAGCAACTATACG-3’; reverse: 5’-CAATGGTCGCCTCTTGTAGA-3’) and 36B4 (forward: 5’-TGCATCAGTACCCCATTCTATCAT-3’; reverse: 5’-AGGCAGATGGATCAGCCAAGA-3’).

### Colony-Forming Assay

Cells (5000–10,000) were plated on 60-mm dishes and incubated at 37°C, 5% CO_2_ for colony formation. After 10 days, colonies were fixed with 10% (v/v) methanol for 15 min and stained with 5% Giemsa (Sigma, St. Louis, USA) for 30 min for colony visualization.

### Respirometry and Extracellular Acidification

The cellular oxygen consumption rate (OCR) and extracellular acidification rate (ECAR) were obtained using a Seahorse XF24 extracellular flux analyzer from Seahorse Bioscience (North Billerica, MA, USA). Experiments were performed according to the manufacturer’s instructions. OCR and ECAR were measured in real time in normoxia, physioxia or hypoxia. 40,000 cells were deprived of glucose for 1 h, then glucose (G–10 mM), oligomycin (O– 1µM), 2,4-Dinitrophenol (DNP–100 µM), and Rotenone + Antimycin A (R/A–1µM) were injected at the indicated times.

### Glucose and Lactate Measurements

The Glucose and lactate concentrations in the supernatant of cells incubated in Hx for 72h was determined by YSI Biochemistry Analyzer. Each condition was determined for 100,000 cells to express the Glucose/Lactate concentration as g/L per 100,00 cells.

### Immunoblotting

Cells were lysed in 1.5x SDS buffer and the protein concentration determined using the BCA assay. 40µg of protein from whole cell extracts were resolved by SDS-PAGE and transferred onto a PVDF membrane (Millipore, Molsheim, France). Membranes were blocked in 5% non-fat milk in TN buffer (50mMTris-HClpH7.4, 150mMNaCl) and incubated in the presence of the primary and then secondary antibodies in 5% non-fat milk in TN buffer. Rabbit polyclonal anti-HIF-1α antibody (antiserum 2087) was produced and characterized in our laboratory (PMID: **10551817**). The antibody against HIF-2α (NB100-122) was purchased from Novus Biologicals (Littleton, CA). ECL signals were normalized to either β-tubulin or HSP90. After washing in TN buffer containing 1% Triton-X100 and then in TN buffer, immunoreactive bands were visualized with the ECL system (Amersham Biosciences, Buckinghamshire, UK).

### Immunocytochemistry

Cells were fixed in 3% paraformaldehyde and extracted with Triton X-100. Primary antibodies included rabbit anti-HIF-1α (PMID: **10551817**) (1:400). Alexa Fluor 594- and 488-conjugated secondary anti-rabbit antibodies (Molecular Probes, Carlsbad, CA, USA) were used at 1:400. Cells were visualized by wide-field, fluorescence microscopy using a DM5500B upright stand (Leica, Wetzlar, Germany) with a 40x oil objective NA 1.00. The cubes used were A4 (excitation filter BP 360/40, dichroic mirror 400, emission filter BP 470/40), L5 (BP 480/40, 05, BP 527/30), and TX2 (BP 560/40, 595, BP645/75). Acquisitions were done with an Orca-ER camera (Hamamatsu, Hamamatsu, Japan). Cells were also visualized using the confocal microscope, Nikon A1R inverted stand (Nikon, Tokyo, Japan). Objectives 10x dry NA 0.3 and/or 40x oil 1.3 NA and/or 60x oil 1.4 NA were used. The lasers used were 405nm, and/or 488nm, and/or 561nm. The microscope was equipped with an automated xy stage for mosaic acquisitions.

### Single Cell Data Availability

The RNA-Seq data discussed in B. Manoranjan’s publication (PMID: **32859895**) have been deposited in NCBI’s Gene Expression Omnibus and are accessible through GEO Series accession number GSE131473. The scRNA-seq data have been deposited in CReSCENT (https://crescent.cloud/; CRES-P22) and processed data is uploaded on the Broad Institute Single Cell Portal https://singlecell.broadinstitute.org/single_cell/study/SCP840.

### Data sources

Primary medulloblastoma RNA-seq data were obtained from the ‘R2: Genomics Analysis and Visualization Platform (http://r2.amc.nl)’ in the data set ‘Tumor Medulloblastoma - Pfister - 223 - MAS5.0 - u133p2’.

Affymetrix Human Gene 1.1 ST Array profiling of 763 primary medulloblastoma samples used for identification of Medulloblastoma subtypes^7^.

### Statistics

Statistical analysis of all data was performed using GraphPad Prism software, version 9.0 (GraphPad Software, La Jolla, CA, USA) and expressed as means ± S.E.M. For multiple comparisons, two-way ANOVA (post hoc Bonferroni) was done. The p-values are indicated (p<0.05 *, p<0.005 **, p<0.0005 ***, p<0.0001 ****) and p-values between 0.05 and 0.10 indicated a statistical tendency.

## Notes

### Competing Interest Statement

The authors have declared no competing interest.

## Reference

1 Petrova, V., Annicchiarico-Petruzzelli, M., Melino, G. & Amelio, I. The hypoxic tumour microenvironment. Oncogenesis 7, 10 (2018). 10.1038/s41389-017-0011-9

2 Pouyssegur, J., Dayan, F. & Mazure, N. M. Hypoxia signalling in cancer and approaches to enforce tumour regression. Nature 441, 437–443 (2006). 10.1038/nature04871

3 Northcott, P. A. et al. The whole-genome landscape of medulloblastoma subtypes. Nature 547, 311–317 (2017). 10.1038/nature22973

4 Northcott, P. A. et al. Medulloblastomics: the end of the beginning. Nat Rev Cancer 12, 818–834 (2012). 10.1038/nrc3410

5 Northcott, P. A. et al. Rapid, reliable, and reproducible molecular sub-grouping of clinical medulloblastoma samples. Acta Neuropathol 123, 615–626 (2012). 10.1007/s00401-011-0899-7

6 Taylor, M. D. et al. Molecular subgroups of medulloblastoma: the current consensus. Acta Neuropathol 123, 465–472 (2012). 10.1007/s00401-011-0922-z

7 Cavalli, F. M. G. et al. Intertumoral Heterogeneity within Medulloblastoma Subgroups. Cancer Cell 31, 737–754 e736 (2017). 10.1016/j.ccell.2017.05.005

8 Manoranjan, B., Adile, A. A., Venugopal, C. & Singh, S. K. WNT: an unexpected tumor suppressor in medulloblastoma. Mol Cell Oncol 7, 1834903 (2020). 10.1080/23723556.2020.1834903

9 Lim, J. H. et al. Bafilomycin induces the p21-mediated growth inhibition of cancer cells under hypoxic conditions by expressing hypoxia-inducible factor-1alpha. Mol Pharmacol 70, 1856–1865 (2006). 10.1124/mol.106.028076

10 Han, D. et al. Lysine methylation of transcription factors in cancer. Cell Death Dis 10, 290 (2019). 10.1038/s41419-019-1524-2

11 Bao, L. et al. Methylation of hypoxia-inducible factor (HIF)-1alpha by G9a/GLP inhibits HIF-1 transcriptional activity and cell migration. Nucleic acids research 46, 6576–6591 (2018). 10.1093/nar/gky449

12 Ballabio, C. et al. Modeling medulloblastoma in vivo and with human cerebellar organoids. Nat Commun 11, 583 (2020). 10.1038/s41467-019-13989-3

13 Schodel, J. et al. Hypoxia, Hypoxia-inducible Transcription Factors, and Renal Cancer. Eur Urol 69, 646–657 (2016). 10.1016/j.eururo.2015.08.007

14 Koh, M. Y. & Powis, G. Passing the baton: the HIF switch. Trends Biochem Sci 37, 364–372 (2012). 10.1016/j.tibs.2012.06.004

15 Pei, Y. et al. An animal model of MYC-driven medulloblastoma. Cancer Cell 21, 155–167 (2012). 10.1016/j.ccr.2011.12.021

16 Das, B. et al. MYC Regulates the HIF2alpha Stemness Pathway via Nanog and Sox2 to Maintain Self-Renewal in Cancer Stem Cells versus Non-Stem Cancer Cells. Cancer Res 79, 4015–4025 (2019). 10.1158/0008-5472.CAN-18-2847

17 Covello, K. L. et al. HIF-2alpha regulates Oct-4: effects of hypoxia on stem cell function, embryonic development, and tumor growth. Genes Dev 20, 557–570 (2006). 10.1101/gad.1399906

18 Petruzzelli, R., Christensen, D. R., Parry, K. L., Sanchez-Elsner, T. & Houghton, F. D. HIF-2alpha regulates NANOG expression in human embryonic stem cells following hypoxia and reoxygenation through the interaction with an Oct-Sox cis regulatory element. PLoS One 9, e108309 (2014). 10.1371/journal.pone.0108309

19 Simon, M. C. & Keith, B. The role of oxygen availability in embryonic development and stem cell function. Nat Rev Mol Cell Biol 9, 285–296 (2008). 10.1038/nrm2354

20 Northcott, P. A. et al. Medulloblastoma. Nat Rev Dis Primers 5, 11 (2019). 10.1038/s41572-019-0063-6

21 Garancher, A. et al. NRL and CRX Define Photoreceptor Identity and Reveal Subgroup-Specific Dependencies in Medulloblastoma. Cancer Cell 33, 435–449 e436 (2018). 10.1016/j.ccell.2018.02.006

